# Tokenvizz: GraphRAG-Inspired Tokenization Tool for Genomic Data Discovery and Visualization

**DOI:** 10.1101/2024.12.03.626631

**Authors:** Çerağ Oğuztüzün, Zhenxiang Gao, Rong Xu

## Abstract

**Summary:** One of the primary challenges in biomedical research is the interpretation of complex genomic relationships and the prediction of functional interactions across the genome. Tokenvizz is a novel tool for genomic analysis that enhances data discovery and visualization by combining GraphRAG-inspired tokenization with graph-based modeling. In Tokenvizz, genomic sequences are represented as graphs, where sequence k-mers (tokens) serve as nodes and attention scores as edge weights, enabling researchers to visually interpret complex, non-linear relationships within DNA sequences. Through a web-based visualization interface, researchers can interactively explore these genomic relationships and extract biologically meaningful insights about regulatory patterns and functional elements. Applied to promoter-enhancer interaction prediction tasks, Tokenvizz outperformed traditional sequential models while providing interpretable insights into genomic features, demonstrating the advantage of graph-based representations for biological discovery.

**Availability and Implementation:** Tokenvizz, along with its user guide, is freely accessible on GitHub at: https://github.com/ceragoguztuzun/tokenvizz.

**ACM Reference Format:** Çerağ Oğuztüzün, Zhenxiang Gao, and Rong Xu. 2024. Tokenvizz: GraphRAG Inspired Tokenization Tool for Genomic Data Discovery and Visualization. In *Proceedings of (Bioinformatics)*. ACM, New York, NY, USA, 7 pages. https://doi.org/XXXXXXX.XXXXXXX

## 1 INTRODUCTION

Understanding how genes interact and regulate biological processes remains a fundamental challenge in biomedical research, with implications ranging from disease treatment to personalized medicine. Modern genomic studies generate vast amounts of complex data about DNA sequences and their functional elements, such as enhancers and promoters that control gene expression [1]. However, making sense of these intricate relationships and extracting meaningful biological insights from genomic data continues to challenge researchers. The ability to interpret these relationships is crucial for identifying key genetic features that drive diseases, understanding how variations in DNA affect biological functions, and ultimately developing targeted therapeutic strategies. While traditional sequential approaches and large language models (LLMs) have proven valuable for processing genomic information, they often struggle to capture and explain the complex, non-linear relationships between genomic elements in an intuitive way [2, 3]. Graph-based models offer a promising alternative by providing a visual and intuitive framework for representing these relationships, enhancing our ability to understand complex genomic interactions and trace the mechanisms underlying genetic variation and disease progression [4–6]

Retrieval-Augmented Generation (RAG) is a technique used to improve LLM outputs by leveraging supplementary information [7]. GraphRAG builds on this by utilizing LLMs to create a knowledge graph from an input corpus. This approach involves slicing the input corpus into smaller units, called TextUnits [7], which are then analyzed to provide ne-grained references. Using an LLM, GraphRAG extracts entities, relationships, and key claims from these TextUnits, and creates a graph, allowing the model to generate more structured and context-rich outputs.

Previous approaches to modeling DNA sequences as graphs faced significant limitations in handling the scale and complexity of genomic data. Early work by [8] and assembly approaches reviewed by [9] struggled to effectively process the massive volume of genomic data while maintaining biological relevance. While Blazewicz et al. traced the evolution from simple Hamiltonian and Eulerian path-based models to more complex overlap and decomposition-based graphs, these traditional approaches focused primarily on assembly rather than relationship discovery. In earlier work like [8], each DNA sequence was represented as a weighted directed multi-graph where nodes were individual nucleotides, and edges were weighted based on metrics like Euclidean distance, cosine similarity, or correlation coefficients. However, the node embeddings used at that time were less informative, as they did not benefit from modern attention mechanisms. Using attention-based architectures, we are able to capture more context-aware representations. Recent work in graph-based genomic analysis includes GraphDSB [10], which demonstrated the effectiveness of graph neural networks for predicting double-strand DNA breaks. While GraphDSB successfully utilized structural encodings and attention mechanisms, it was limited by its specialized focus and fixed graph construction using 5-kb genome bins. Tokenvizz addresses these limitations by providing a flexible, general-purpose framework that leverages pre-trained genomic language models and customizable tokenization strategies. Moreover, the GraphRAG approach has been applied to medical datasets to improve answering medical questions, providing evidence-based responses, and enhancing safety and reliability [11]. However, it has yet to be used for processing DNA sequences.

We propose the Tokenvizz tool, a genomic sequence tokenizer and graph modeling approach inspired by GraphRAG. It is a discovery tool and also features a web-based visualizer that allows researchers to intuitively explore relationships between different parts of a DNA sequence. Tokenvizz tokenizes the input genomic sequence, represents each token as a node in a graph, and uses attention scores - weights that capture the task-specific importance and relationships between tokens [12] - to dene edge weights, thereby revealing patterns within the sequence. To demonstrate Tokenvizz’s utility, we applied it to promoter-enhancer interaction prediction which is a critical task in understanding gene regulation. Furthermore, the graph representations generated by Tokenvizz can be used as inputs for other models, and Tokenvizz is built to be extensible, allowing users to integrate additional layers of information into the graph, such as various node and edge types that represent domain-specific data relevant to their analyses.

## 2 METHODS AND IMPLEMENTATION

Tokenizz is composed of four main modules: (i) data processing, (ii) tokenization, (iii) graph construction, and (iv) visualization (Figure 1).

**Figure 1:**
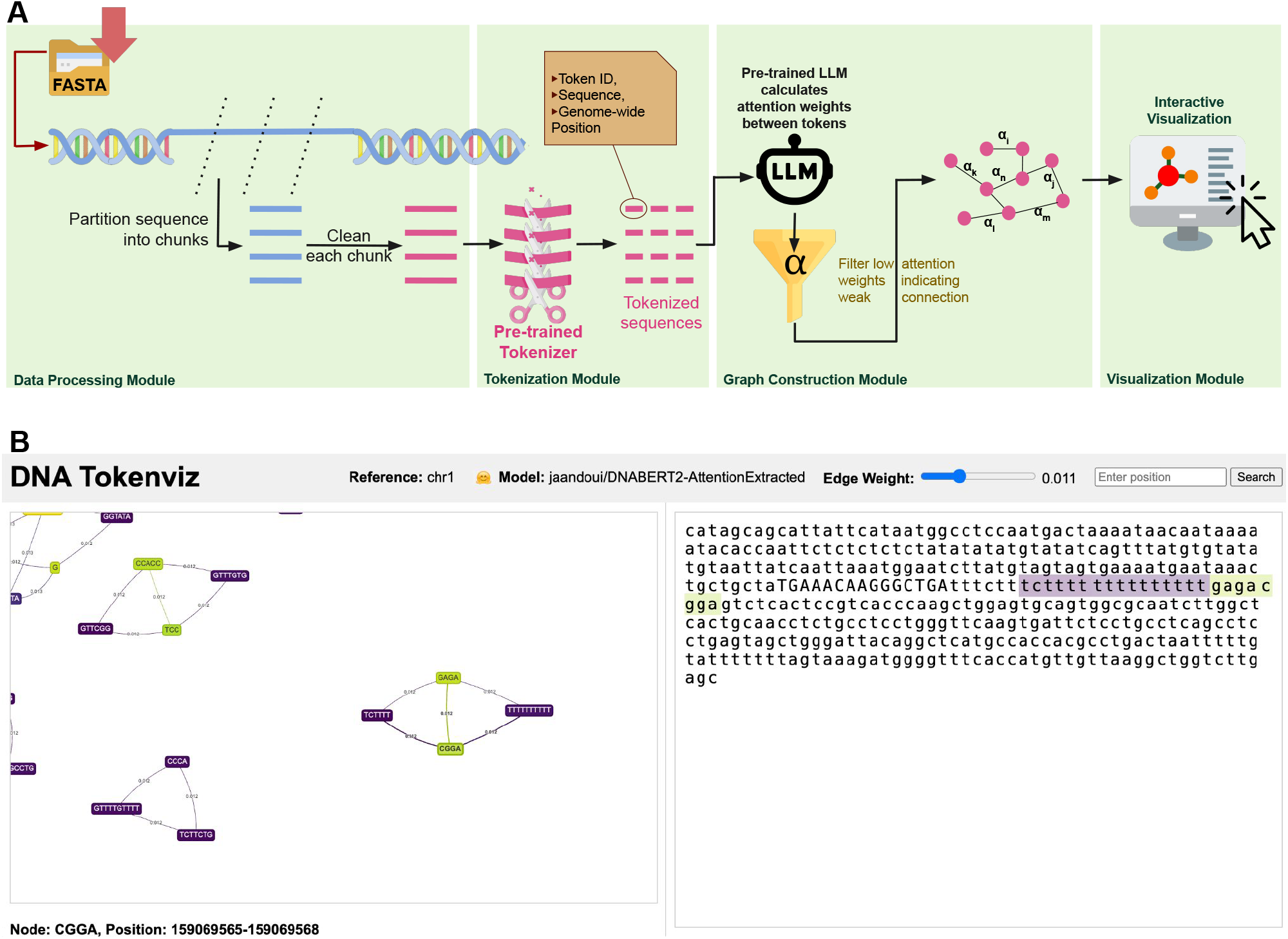
(A) Overview of Tokeniz. The user inputs genomic sequences in FASTA format for analysis. The tool processes these sequences using pre-trained tokenizers and language models from Hugging Face selected by the user, to generate a graph representation which is then rendered for interactive visualization. (B) The web-based visualization interface enables dynamic sequence exploration through multiple features. Users can adjust edge weight thresholds to focus on strongly connected subgraphs and perform position-based sequence searches. When clicking a specific node, the interface displays the surrounding sequence context with highlighted clicked node and neighbors, while also showing the genome-wide position and detailed sequence information in the lower panel.

### 2.1. Data Processing Module

We designed the data preprocessing module for Tokenvizz to efficiently handle large genomic sequences and prepare them for processing by the genomic language models (gLMs). The pseudocode of this module is provided in Algorithm 1 in Supplementary. From the FASTA le inputted by the user (we used the human reference genome hg38), we extracted references (such as chromosomes) and sequence lengths to systematically iterate over them. The module divides each large DNA sequence into calculated chunks, whose sizes are based on the model’s maximum sequence length, adjusting for any special tokens required by the tokenizer. This ensures that each subsequence fits within the model’s input limits. The chunking process accounts for sequences that aren’t perfectly divisible by the chunk size, thereby covering the entire sequence without omission.

We clean each chunk by standardizing nucleotide representation by converting all characters to uppercase and retaining only valid nucleotides (A, T, C, G) using a translation table. Other nucleotides are removed during sequence cleaning, however they are still counted to calculate positions. To handle metadata preservation during batching, we designed a custom collate function that maintains the alignment between sequences and their associated metadata—such as reference names and start positions—when creating batches for the model. To conserve computational resources, each reference in the input FASTA file (e.g., chromosomes in hg38) is represented as a separate graph. Users interested in interchromosomal interactions can input merged sequences into the tool.

### 2.2 Tokenization Module

gLMs utilize various tokenization strategies to effectively process DNA sequences, with the choice of tokenizer significantly impacting model performance and interpretability [13]. Key tokenization approaches include single nucleotide tokenization, where each nucleotide is treated as a separate token; k-mer tokenization; and byte-pair encoding (BPE) [14]. We opted not to implement single nucleotide or overlapping k-mer tokenization due to their computational intensity. Instead, Tokenvizz uses BPE from the DNABERT2 [15] model and non-overlapping k-mer tokenization from the NT model [16]. Additionally, Tokenvizz allows users to select any tokenizer from pre-trained models available on Hugging Face [17] by simply specifying the model’s name.

During tokenization, we configured the tokenizer to return offset mappings, which provide the start and end indices of each token within the original sequence. After tokenization, we iterated over each sequence in the batch, extracting tokens and their associated offset mappings. For each token, we adjusted its positional indices by adding the offset of the sequence chunk within the entire genome, effectively translating chunk-local positions to genome-wide positions.

### 2.3 Graph Construction Module

The token information, including token IDs, token strings, and adjusted genomic positions, was compiled into a structured format for each sequence, denoting a node in our graph. We processed each sequence through the pre-trained gLM to extract attention weights. After obtaining the raw attention weights from the model, we applied a softmax function to these weights to normalize, converting them into probabilities, that allows for a meaningful interpretation of the attention as the strength of relationships between tokens. We averaged the attention scores over all layers and heads of the model. This process is outlined in Algorithm 2 in Supplementary 8.1. To avoid redundant edges and ensure the graph is undirected, we utilized only the upper triangular portion of the attention matrix when establishing connections between nodes. This design choice prevents duplicate edges between the same pairs of nodes.

We developed a method to construct a graph *G*=(*V, E*)where each node *υ*_*i*_ ∈ *V* represents a token as: *υ*_*i*_ = {token_id_*i*_, token_string_*i*_, position_*i*_, and edges *E* {(*υ*_*i*_, *υ*_*j*_) | *a*_*ij*_ ≥ θ and *i* <*j}* where *a*_*ij*_ represents the attention-derived probability from token *υ*_*i*_ to *υ*_*j*_. To focus on meaningful relationships and reduce noise, we implemented threshold filtering on the attention weights, post-processing. Only edges with attention scores above a user-selected threshold value θ are included in the graph. This filtering process excludes weaker connections that could obscure important patterns within the data. After, we also include a degree connectivity threshold (Supplementary, 8.2) to filter out the nodes that have below the number of user-selected threshold of neighbors to ensure strong connectivity in our graph.

### 2.4. Visualization Module

As an additional functionality, we integrated a suite of functionalities to transform processed graph data into an interactive web-based visualization tool. This allows users to visualize the tokenized DNA sequences as dynamic graphs. The frontend of the visualization interface is built using HTML, CSS, and JavaScript.

When users click a node, the tool retrieves the corresponding DNA segment from the genome using a Flask-based backend server. The fetched DNA sequence is then displayed alongside the graph, with the node’s sequence highlighted. Additionally, neighboring nodes are highlighted in the DNA sequence, providing a clear and intuitive connection between the abstract graph representation and the actual sequential genomic representation (Figure 1(B).

### 2.5. Performance

To manage the immense size of genomic data efficiently, we used ‘pysam’ library’s ‘FastaFile’ class [18], which enables random access to specific DNA sequences directly from FASTA files without the need to load the entire genome into memory. Throughout the implementation, we aligned the data preprocessing steps with the model’s requirements by dynamically adjusting chunk sizes based on the model’s maximum sequence length. To leverage parallel computation of GPUs, we employed batch tokenization, which processes multiple sequences simultaneously. We implemented a dedicated data module with Pytorch Lightning [19] which allows adjustments to batch sizes, the number of worker processes, and subsets of data, which is particularly beneficial when scaling the application or conducting tests on smaller datasets. Collectively, these performance-focused implementations ensure that Tokenvizz can effiectively process the whole human genome of 3b base pairs.

In this workflow, a DNA sequence of length *n* is tokenized into *m* tokens, where *m* is linearly proportional to = (i.e., *m* = O(*n*)). The runtime of Tokenvizz’s workflow is fundamentally O(*n*^2^), due to the self-attention mechanism inherent in the gLM utilized for processing DNA sequences. We have created the graphs for the hg38 using one NVIDIA A100 80GB PCIe GPU, but shorter sequences require less computation.

## 3 APPLICATION

To demonstrate Tokenvizz’s effiectiveness as a genomic analysis tool, we evaluated our graph-based approach on 6 publicly available datasets from the DNABERT2 [15] study. Using the Genome Understanding Evaluation (GUE+) dataset [15], we tested Tokenvizz on enhancer-promoter interaction prediction, a binary classification task predicting interactions between 5,000-nucleotide sequences. We implemented a straightforward Graph Convolutional Network (GCN)-based [20] classification pipeline using Tokenvizz’s graph representations (architecture details in Supplementary). When benchmarked against Matthews Correlation Coefficient (MCC) scores [21] from DNABERT2 [15] and Nucleotide Transformer (NT) [16], our approach consistently outperformed these state-of-the-art models. These results demonstrate that Tokenvizz’s graph-based modeling can capture complex biological interactions, validating its utility as a tool for enhanced genomic analysis. Building from this application, we provide an example demonstrating Tokenvizz’s interpretability in Supplementary Section 8.4.

**Table 1:** Matthews Correlation Coefficient (MCC) for different models across datasets for enhancer-promoter interaction prediction task.

| Datasets | DNABERT2 | NT | Tokenvizz (BPE) |
| --- | --- | --- | --- |
| NHEK | 76.21 | 61.91 | **82.55** |
| K562 | 79.19 | 72.15 | **85.07** |
| IMR90 | 83.50 | 73.13 | **84.27** |
| HeLaS3 | 86.71 | 79.49 | **91.85** |
| HUVEC | 92.90 | 86.48 | **95.46** |
| GM12878 | 73.70 | 68.64 | **76.57** |

Note: 'BPE' indicates that Tokenvizz used DNABERT2's BPE tokenizer [15] to create the graph model.

## 4 CONCLUSION

We present Tokenvizz, a novel tool for enhanced genomic sequence analysis through attention-based graph modeling, also featuring interactive visualization capabilities to support data exploration. By tokenizing genomic sequences into k-mer nodes and utilizing attention scores to map their relationships, Tokenvizz offers a framework for understanding complex genetic interactions. Our application to promoter-enhancer prediction demonstrates superior performance over traditional sequential methods. The tool’s extensible architecture enables integration of diverse biological data sources, while its intuitive web-based interface makes complex genomic analysis accessible to researchers. Looking ahead, we plan to expand Tokenvizz’s capabilities through integration with downstream applications in predictive modeling and functional genomics.

## ACKNOWLEDGMENTS

This work has been supported by NIH National Institute of Aging, USA R01 AG057557, R01 AG061388, R56 AG062272, National Institute on Alcohol Abuse and Alcoholism, USA (grant no. R01AA029831), National Eye Institute, USA (EY029297), National Institute on Drug Abuse, USA (UG1DA049435, CTN-0114), American Cancer Society Research Scholar, USA Grant RSG-16-049-01-MPC, The Clinical and Translational Science Collaborative (CTSC) of Cleveland, USA (UL1TR002548-01).

## 6. DECLARATIONS

The authors declare no competing interests. Ethics approval and consent to participate is not applicable.

## 7 DATA AVAILABILITY

The data used in *Section 3. Application* is freely accessible in the DNABERT2 manuscript [15], which also provides additional information about the datasets.

## 8 CODE AVAILABILITY

The Tokenvizz tool is freely available on GitHub with a user guide at https://github.com/ceragoguztuzun/tokenvizz.

