## Supplementary Materials for "Tokenvizz: GraphRAG-Inspired Tokenization Tool for Genomic Data Discovery and Visualization"

### 8.1 Algorithm Pseudocodes

**Algorithm 1** Data Pre-processing for Tokenvizz

**Require:** `fasta_file_path`: Path to the input FASTA file (e.g., hg38)
**Require:** `model`: Pre-trained genomic Language Model (gLM)
**Ensure:** Preprocessed DNA sequence chunks with associated metadata

**Parameters:**
  $M \leftarrow model$.max_seq_length ▷ Maximum sequence length for the model
  $S \leftarrow model$.tokenizer.num_special_tokens_to_add(pair=False) ▷ Number of special tokens required by tokenizer
  $C \leftarrow M - S$ ▷ Chunk size
**Load FASTA File:**
  `fasta` ← `pysam.FastaFile(fasta_file_path)`
  `references` ← `fasta.references`
  `lengths` ← `fasta.lengths`
**for** `i` **in** `range(len(references))` **do**
    `ref` ← `references[i]`
    $L_i \leftarrow$ `lengths[i]` ▷ Length of chromosome `ref`
    $n_i \leftarrow \lceil \frac{L_i}{C} \rceil$ ▷ Number of chunks for `ref`
  **for** `j` **in** `range(n_i)` **do**
      $start \leftarrow j \times C$
      $end \leftarrow \min((j+1) \times C, L_i)$
      `chunk` ← `fasta.fetch(ref, start, end)`
      **Clean the Chunk:**
        `clean_chunk` ← ”.join([base for base in chunk.upper() if base in {'A', 'T', 'C', 'G'}])
      **Preserve Metadata:**
        `metadata` ← { 'reference': `ref`, 'start': *start*, 'end': *end* }
      **Process the Chunk:**
        `process_chunk(clean_chunk, metadata)` ▷ involves further steps such as tokenization and mapping.
    **end for**
**end for**

### 8.2 Weighted Degree Centrality Formulation

The Weighted Degree Centrality $C_i$ of a node $v_i \in V'$ quantifies the total strength of connections that $v_i$ has with other nodes in the graph. It is calculated as the sum of the weights of all edges incident to $v_i$.

$$C_i = \sum_{v_j \in N(v_i)} w_{ij} \tag{1}$$

where $N(v_i)$ is the set of neighbors of node $v_i$ in the graph $G' = (V', E')$. $w_{ij}$ is the weight of the edge $(v_i, v_j)$, corresponding to the attention-derived probability $a_{ij}$.

**Algorithm 2** Calculate Attention Weights for Tokenvizz

**Require:** $\mathcal{A}$: Batch of attention weight tensors for each sequence
**Ensure:** $\bar{\mathcal{A}}$: List of averaged attention matrices for each sequence
Initialize $\bar{\mathcal{A}} \leftarrow []$
**for** $i \in \{1, 2, \ldots, |\mathcal{A}|\}$ **do** ▷ Each sequence in the batch
  $\mathbf{A}_i \leftarrow \mathcal{A}[i]$ ▷ Attention weights for sequence $i$
  Initialize $\tilde{\mathbf{A}}_i \leftarrow []$
  **for** $l \in \{1, 2, \ldots, L\}$ **do** ▷ Iterate over layers
    **for** $h \in \{1, 2, \ldots, H\}$ **do** ▷ Iterate over heads
      $\tilde{\mathbf{A}}_i^{(l,h)} \leftarrow \text{softmax}(\mathbf{A}_i^{(l,h)}, \text{dim} = -1)$ ▷ Normalize attention weights
      $\tilde{\mathbf{A}}_i \leftarrow \tilde{\mathbf{A}}_i \cup \{\tilde{\mathbf{A}}_i^{(l,h)}\}$ ▷ Store in list
    **end for**
  **end for**
  $\bar{\mathbf{A}}_i \leftarrow \frac{1}{L \times H} \sum_{l=1}^{L} \sum_{h=1}^{H} \tilde{\mathbf{A}}_i^{(l,h)}$ ▷ Average over layers and heads
  $\bar{\mathcal{A}} \leftarrow \bar{\mathcal{A}} \cup \{\bar{\mathbf{A}}_i\}$
**end for**
**return** $\bar{\mathcal{A}}$ ▷ Contains the averaged attention matrices for each sequence

### 8.3 Model Architecture and Hyperparameter Optimization for Application

The study employs a Graph Convolutional Network (GCN) [20] architecture tailored for genomic sequence classification. The model consists of 3 linear layers with hidden channel sizes of 128 and 256, each followed by batch normalization and a dropout rate of 0.3 to prevent overfitting. Utilizing the AdamW optimizer [22], the network is trained for 300 epochs with a weight decay of 1e-6 to enhance generalization. Hyperparameters were systematically selected through `GridSearchCV` [23] to optimize the Matthews Correlation Coefficient (MCC) [21] across 3-fold cross-validation. The model captures complex k-mer interactions within genomic graphs by leveraging class-weighted binary cross-entropy loss [24].

### 8.4 Example for Interpretability

In our application, we classified enhancer-promoter sequence pairs from the GUE+ HUVEC dataset [15] using Tokenvizz. To illustrate interpretability, we randomly selected a sequence pair from the test set that was both labeled and predicted as interacting. We then concatenated the sequences with a separator to form a single continuous sequence, which we used to generate a graph using Tokenvizz (Figure 2).

We observed core promoter elements (TATA box, BREu, etc.) in the *promoter region* and identified their corresponding nodes and weighted degrees. Tokenvizz successfully identified BREu motifs with high weighted degrees (0.98-0.99) which indicates their functional importance (a single example is provided in Figure 3).

For the *enhancer region*, we performed Louvain community detection [25] and revealed several distinct communities with high modularity (0.9275), each containing biologically relevant motif patterns. These communities show varying levels of connectivity and clustering (coefficients ranging from 0.1889 to 0.6723), which were

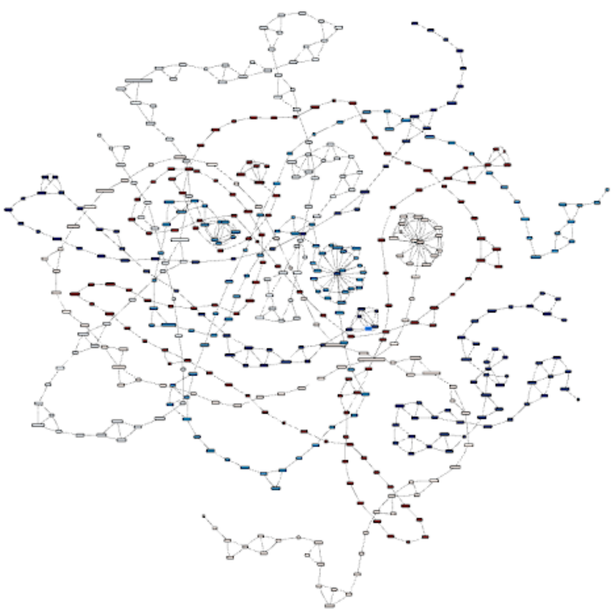

Figure 2: The graph visualizes enhancer nodes (blue) and promoter nodes (red) for an example application. Deeper color signifies nodes with higher weighted degree centrality. Tokenvizz automatically separates enhancers (blue) and promoters (red) into distinct subgraphs, reflecting the biological modularity of regulatory elements.

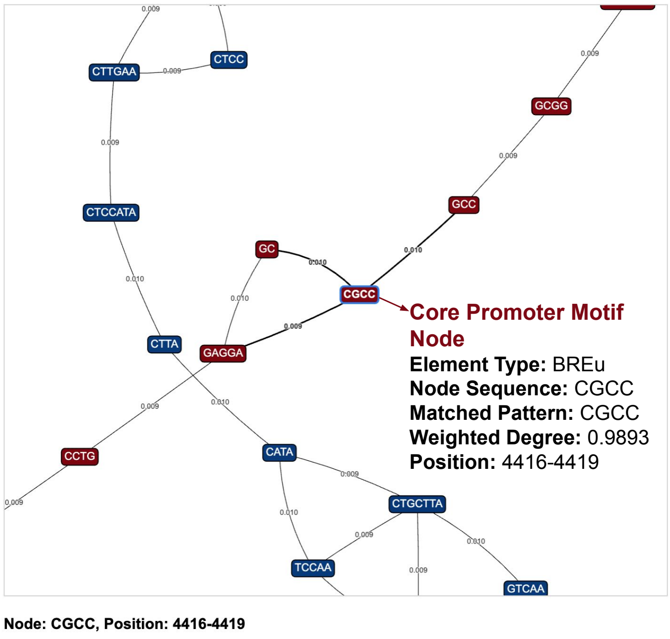


Figure 3: Visualization of the BREu motif node within the promoter region identified by Tokenvizz. This high weighted degree of the node (0.9893) indicates the functional importance of the BREu motif in gene regulation.

validated using MAST motif discovery tool [26] (a single example is provided in Figure 4).

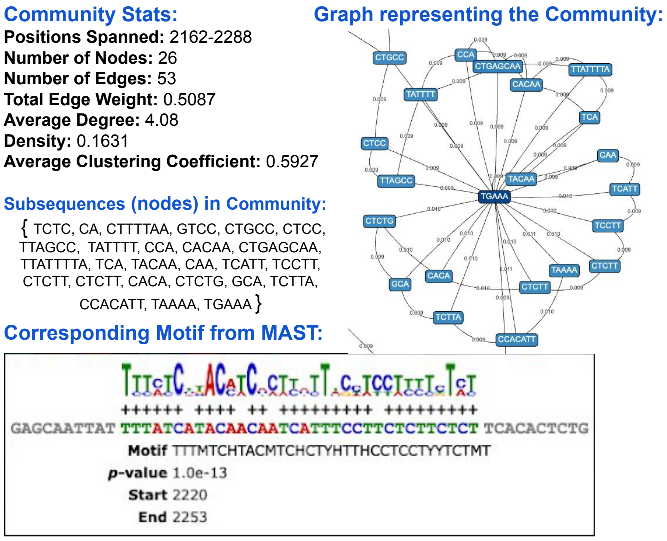


Figure 4: The subgraph representing a community within the enhancer sequence reflects the motif's patterns, with its nodes corresponding to the motif's subsequences.

Tokenvizz provides a more intuitive view of complex regulatory relationships through its graph structure. Network metrics such as density, clustering coefficients, and edge weights offer quantifiable insights that can be compared with known biological patterns. This combination of visual representation and measurable outputs may make Tokenvizz's results more interpretable and aligned with biological understanding than the attention scores of sequential models.
